# Sex-dependent Relationship Between Wrist Deviation and Scaphoid Kinematics

**DOI:** 10.1101/298398

**Authors:** Marissa Borgese, Brent Foster, Robert D. Boutin, Christopher O. Bayne, Robert M. Szabo, Abhijit J. Chaudhari

## Abstract

Several methods of describing patterns of carpal kinematics from radiographs have emerged due to their potential use in developing personalized treatments for wrist pathologies. Such radiography-derived metrics have been used to infer associations between patterns of scaphoid kinematics and other clinically relevant factors such as sex, but the simultaneous effects of sex and wrist position on scaphoid kinematic metrics has yet to be considered. We sought to investigate the relationship between wrist position in the coronal plane and radiographic measurements of the scaphoid for each sex independently, then identify sex-specific differences in scaphoid measurements and calculated metrics. We retrospectively identified 38 subjects with posteroanterior radial and ulnar deviation radiographs. Radiographic scaphoid measurements were collected and used to calculate five scaphoid kinematic metrics per participant. We used Pearson correlation coefficients to explore the relationships between the degrees of wrist deviation in the coronal plane and radiographic scaphoid measurements for men and women independently. We used the non-parametric Wilcoxon signed-rank test to compare values between sexes. The correlations between degrees of coronal wrist deviation and radiographic measurements of scaphoid inclination were significant only for men. Men also had significantly greater values for all radiographic scaphoid measurements. Our study demonstrated sex-specific differences in the relationship between the degrees of radial and ulnar wrist deviation and scaphoid positioning.

**Clinical Relevance:** Our findings show the importance of stratifying by sex in studies of carpal kinematics, such as scaphoid kinematics, and that investigation of strategies to restore normal carpal function should incorporate sex as a biological variable.

## INTRODUCTION

Understanding the baseline kinematic behaviors of the carpal bones in the wrist can aid in the development of personalized treatments for wrist pathologies. Thus several methods of describing the existing variation in carpal kinematic patterns have emerged. Theories of carpal kinematics available in the literature broadly fall within the spectrum of carpal movement described by the ‘row-column’ theory presented by Craigen and Stanley^1^. This theory describes the trajectory of the scaphoid bone on radiographs from radial to ulnar deviation of the wrist, where ‘row’ wrists exhibit movement along the path of radial/ulnar deviation (Fig 1) and ‘column’ wrists display more flexion/extension movement within the proximal carpal row (Fig 2). Studies have since elaborated on the work of Craigen and Stanley^1^ by proposing additional radiography-derived measurements and calculated metrics of scaphoid kinematics that incorporate changes in scaphoid position and length with radial and ulnar wrist deviation^2,3^. Radiographic metrics provide quantitative data for evaluating associations between patterns of scaphoid kinematics and other clinically relevant factors. Prior studies of scaphoid kinematic patterns have revealed associations with both ligamentous laxity^2^ and sex^1^, where women and individuals with more lax wrists are expected to have ‘column’ wrists and men are expected to have ‘row’ wrists. However, the effect of the degrees of radial and ulnar wrist deviation at the time of radiograph acquisition have yet to be considered. Given the known relationship between laxity and sex^2^ and the indirect association of radial and ulnar wrist deviation with wrist laxity^4^, wrist position and patient sex should simultaneously affect scaphoid kinematics.

**Figure 1.**
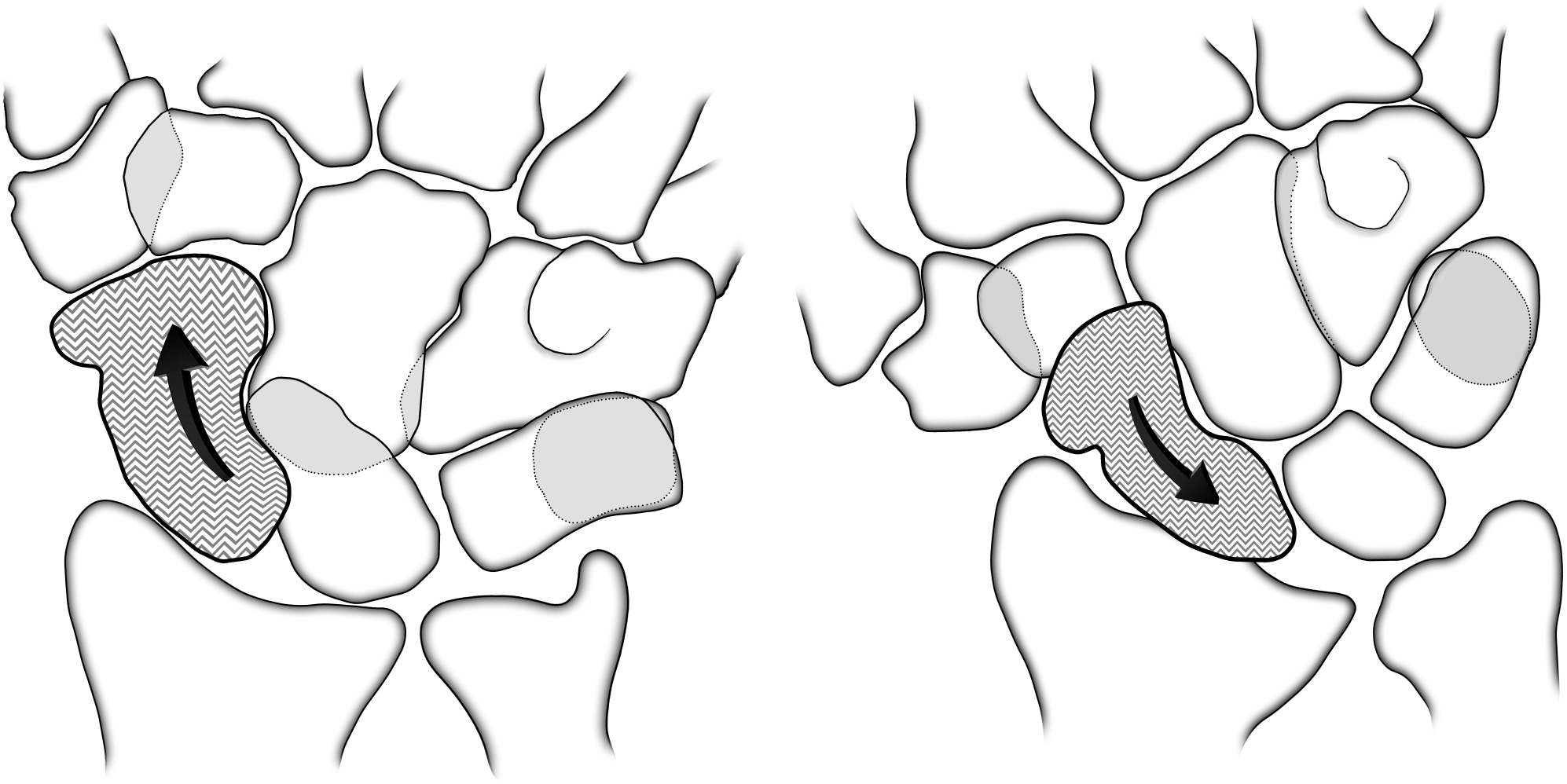
Position of the scaphoid during ulnar and radial deviation of a ‘row’ wrist

**Figure 2.**
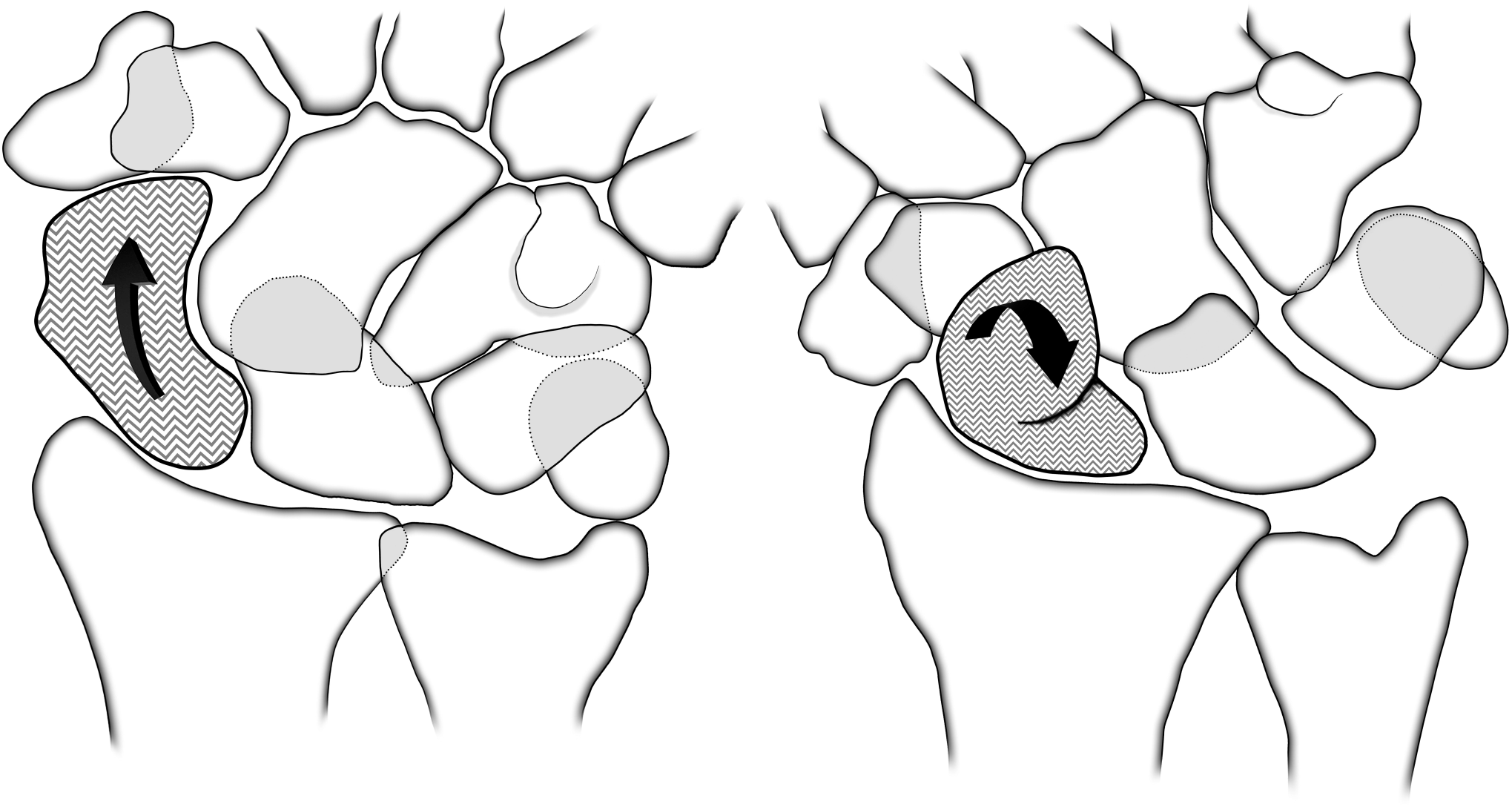
Position of the scaphoid during ulnar and radial deviation of a ‘column’ wrist

The primary purpose of this study was to investigate the relationship between wrist position in the coronal plane and radiographic measurements of the scaphoid bone for men and women independently. Our second objective was to determine whether the calculated metrics of scaphoid kinematics presented by Craigen and Stanley^1^ and Garcia-Elias et al.^2^ would differ significantly between the sexes despite variation in radial and ulnar deviation wrist positioning.

## METHODS

Our study provides level II evidence using a retrospective cohort of consecutively identified patients.

### Subject selection

Our study received local Institutional Review Board approval and the requirement for informed consent was waived because it was done retrospectively and without collecting patient identifiers. We identified participants from our Department of Radiology Picture Archiving and Communication System database by running a search using keywords ‘wrist’ and ‘MR arthrogram’ for individuals of both sexes in the 18 to 45-year age range who had posteroanterior wrist radiographs in radial and ulnar deviation. We chose the 18 to 45-year age range to reduce potential confounding by age-related wrist disorders. Our inclusion criteria included available MR arthrograms, which were used to exclude patients with lunotriquetral and scapholunate interosseous ligament tears, and posteroanterior radial and ulnar deviation radiographs, which were used to collect wrist deviation and scaphoid measurements. We collected the sex of each patient.

We excluded twenty-three subjects with lunotriquetral interosseous ligament tears, fifteen subjects with scapholunate ligament tears, twelve patients with both lunotriquetral and scapholunate interosseous ligament tears, two subjects with orthopaedic hardware, one with lunotriquetral coalition, and one with Ehlers-Danlos syndrome. Thirty-eight subjects (23 men and 15 women) qualified according to the study inclusion and exclusion criteria. Images were evaluated by a radiology researcher with one year of experience in radiographic assessment after obtaining training from a board certified musculoskeletal radiologist with over 20 years of experience. Measurements were recorded to a tenth of a millimeter using the built-in ruler tool in the Philips Picture Archiving and Communication System.

### Wrist deviation

During acquisition of radial and ulnar deviation posteroanterior radiographs at our institution, patients were told to radially and ulnarly deviate their wrists as far as possible. The extent of wrist deviation was found by measuring the angle in degrees between axes drawn along the radius and along the third metacarpal bone on conventional radiographs acquired in both radial and ulnar wrist deviation. This follows commonly accepted methods from wrist goniometry and radiography^5^.

### Radiographic measurements

We collected two measurements from each radiograph, resulting in four reference radiographic measurements per participant (Fig 3). Scaphoid length (R1, R2) and scaphoid inclination (R3, R4) were measured as proposed by Garcia-Elias et al.^2^. The length of the scaphoid was measured as the distance between the most proximal and ulnar point to the central crest on the distal pole in ulnar (R1) and radial (R2) deviation. Scaphoid inclination was measured as the distance between a vertical line intersecting the most proximal and ulnar point on the scaphoid and the radial-most point on the radial styloid in radial (R3) and ulnar (R4) deviation.

**Figure 3.**
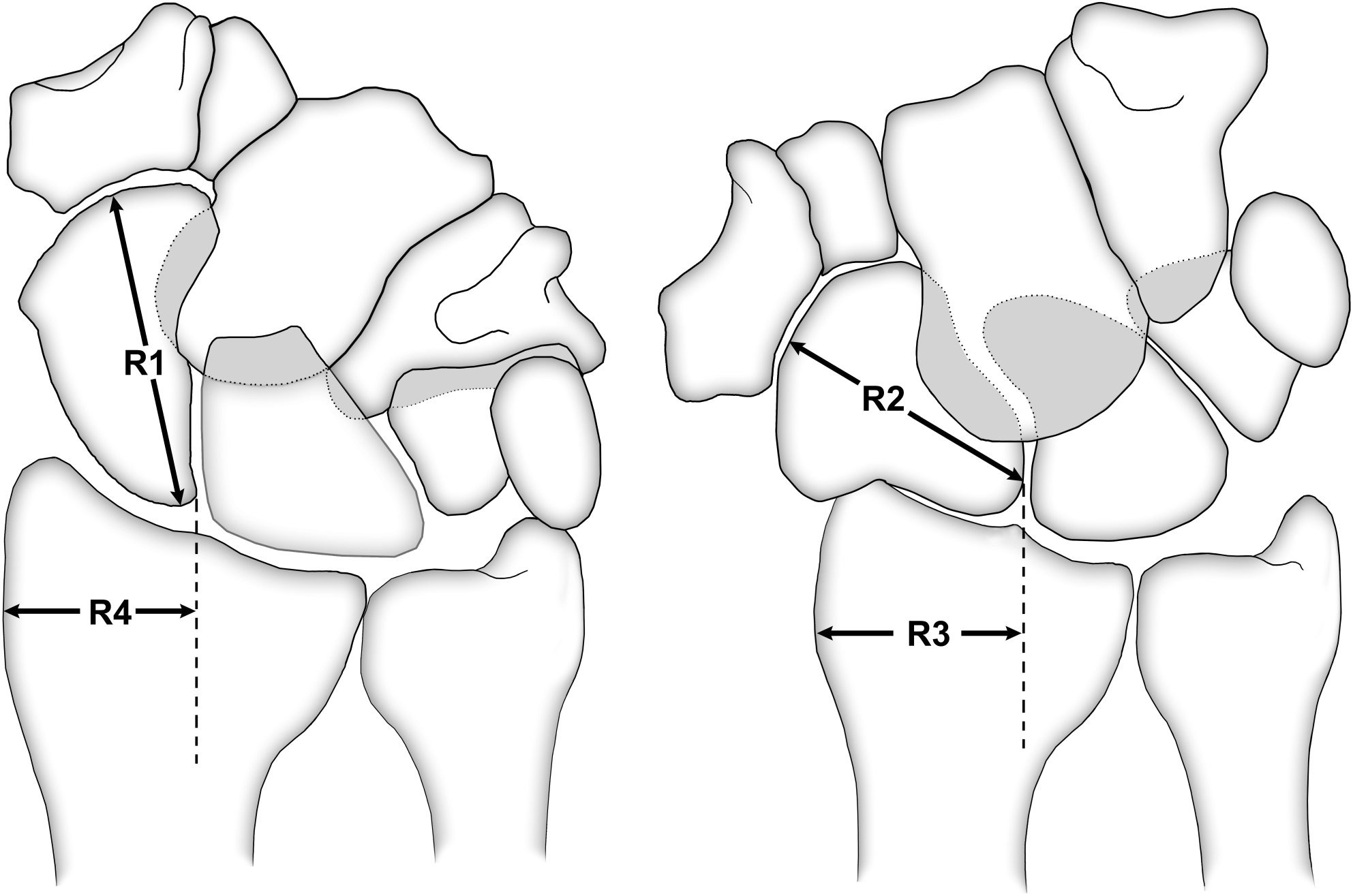
Scaphoid measurements and calculated scaphoid kinematic metrics

### Intra-observer agreement

To assess the intra-observer agreement for each radiographic measurement (R1, R2, R3, R4), a twenty-subject subset of the study cohort was randomly selected and measurements were repeated by the observer one month after the initial measurement collection.

### Kinematic metrics

We used the radiographic scaphoid measurements to calculate five scaphoid kinematic metrics per participant. The Scaphoid Flexion Index (SFI), Scaphoid Inclination Index (SII), and Scaphoid Kinematic Index (SKI) were all calculated as proposed by Garcia-Elias et al.^2^.

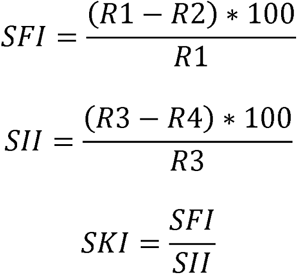

The CR Index (CR) and Translation Ratio (TR) originally proposed by Craigen and Stanley^1^ were calculated using modified equations presented by Galley et al.^3^.

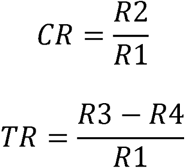

Larger values of the SFI and SKI were meant to identify ‘column’ wrists, while larger values of the SII, CR, and TR were meant to identify ‘row’ wrists.

### Statistical analyses

Due to the non-normal distributions of our data, we used non-parametric Wilcoxon signed-rank tests to compare values of all recorded variables including age, radial and ulnar wrist deviation, and radiographic scaphoid measurements (R1-R4) between the sexes.

We used Pearson correlation coefficients to explore the relationships between the degrees of radial wrist deviation and corresponding radiographic measurements (R2, R3) and between the degrees of ulnar wrist deviation and corresponding radiographic measurements (R1, R4) for men and women independently.

Intra-rater agreements for all four measurements were determined using intraclass correlation coefficients.

We used non-parametric Wilcoxon signed-rank tests to compare values of SFI, SII, SKI, TR, and CR for men versus women. P-values below 0.05 were considered statistically significant for all analyses.

## RESULTS

### Age

Male and female participants did not differ significantly in age (Table 1).

**Table 1.** Median (IQR) is reported due to non-normal data distributions and p-values are for comparisons between men and women using a non-parametric signed-rank test.

|  | Grouped (N=38) | Men (N=23) | Women (N=15) | P-value |
| --- | --- | --- | --- | --- |
| Age (years) | 33 (22-37) | 33 (23-37) | 34 (25-39) | 0.69 |
| Radial deviation (degrees) | 18.0 (13.3-23.0) | 19.0 (14.0-24.5) | 18.0 (13.0-19.0) | 0.60 |
| Ulnar deviation (degrees) | 27.0 (23.0-31.8) | 25.0 (21.5-32.0) | 27.0 (25.0-31.5) | 0.52 |
| R1 (mm) | 23.2 (20.8-25.6) | 24.8 (23.0-26.8) | 20.5 (19.1-23.0) | <0.001 |
| R2 (mm) | 19.5 (17.6-21.6) | 20.8 (19.5-22.2) | 16.9 (14.9-18.5) | <0.001 |
| R3 (mm) | 21.0 (18.6-23.2) | 23.1 (21.9-23.9) | 17.4 (16.8-19.4) | <0.001 |
| R4 (mm) | 17.1 (14.8-18.5) | 18.0 (16.7-20.6) | 14.5 (13.5-16.6) | <0.001 |
| Scaphoid flexion index | 16.5 (12.0-21.2) | 16.1 (11.4-21.1) | 16.9 (13.4-21.7) | 0.44 |
| Scaphoid inclination index | 19.1 (10.3-25.0) | 19.9 (10.4-26.6) | 18.3 (11.1-22.2) | 0.54 |
| Scaphoid kinematic index | 0.88 (0.63-1.66) | 0.82 (0.62-1.11) | 0.97 (0.75-1.72) | 0.30 |
| CR index | 0.84 (0.79-0.88) | 0.84 (0.79-0.89) | 0.83 (0.78-0.87) | 0.44 |
| Translation ratio | 0.16 (0.09-0.22) | 0.19 (0.09-0.26) | 0.15 (0.10-0.21) | 0.50 |

### Wrist deviation

The degrees of ulnar wrist deviation observed in both men and women were generally greater than the degrees of radial wrist deviation observed (Table 1). No significant sex-specific differences in degrees of radial or ulnar deviation were observed.

### Radiographic measurements

Men had significantly greater values of all radiographic scaphoid measurements (R1-R4) compared to women (Table 1).

The correlation between R1 and degrees of ulnar wrist deviation was positive and weak for men and negligible for women (Table 2). The correlation between R2 and degrees of radial wrist deviation was negligible for both sexes. The correlation between R3 and radial wrist deviation was positive and moderate for men and negligible for women. The correlation between R4 and degrees of ulnar wrist deviation was negative and moderate for men and negative and weak for women. Only the correlations of R3 with degrees of radial deviation and R4 with ulnar deviation were statistically significant, and only for men. Medians and interquartile ranges for each radiographic measurement for each sex and with sexes grouped can be found in Table 1. Pearson correlation coefficients and p-values can be found in Table 2.

**Table 2.** Values given are Pearson correlation coefficients (r) and p-values (p).

|  | Men |  | Women |  |
| --- | --- | --- | --- | --- |
| Variables | r | p | r | p |
| R1 vs. Ulnar deviation | 0.26 | 0.22 | 0.08 | 0.78 |
| R2 vs. Radial deviation | 0.05 | 0.81 | 0.13 | 0.63 |
| R3 vs. Radial deviation | 0.45 | 0.03 | 0.13 | 0.63 |
| R4 vs. Ulnar deviation | -0.59 | <0.01 | -0.20 | 0.47 |

### Intra-observer agreement

Intra-rater agreement was excellent for all four radiographic measurements with an ICC of 0.99, 0.85, 0.95, and 0.94 for R1, R2, R3, and R4 respectively.

### Kinematic metrics

Values of SFI, SII, SKI, CR, and TR did not differ significantly for comparisons between men and women. Medians and interquartile ranges of the metrics of scaphoid kinematics for each sex, along with p-values, can be found in Table 1.

## DISCUSSION

Men had significantly greater values of all radiographic scaphoid measurements (R1-R4) compared to women, likely due to the larger overall size of male wrists^6^. We also found sex-related differences in the associations of the degree of wrist deviation with radiographic measurements of scaphoid inclination. The measurement of scaphoid inclination derived from radial deviation wrist radiographs (R3) had a stronger correlation with the degree of radial wrist deviation in men compared to women. The measurement of scaphoid inclination derived from ulnar deviation wrist radiographs (R4) had a stronger correlation with the degree of ulnar wrist deviation in men compared to women.

Based on our results, we accepted our hypothesis and supported the previously reported assertion that men tend to have more ‘row’ wrists^1^. Specifically, we found that measures of scaphoid inclination had significant associations with the degrees of wrist deviation for men only, which would be expected as the proximal scaphoid slides along the distal radius in the frontal plane as the wrist becomes more ulnarly deviated, decreasing the distance between the ulnar-sided scaphoid and the radial-sided radius. While women generally had larger SFI values than men, the difference was not statistically significant. However, women had little change in scaphoid inclination with wrist deviation, suggesting greater scaphoid movement in the sagittal plane and signifying more ‘column’-type wrists.

The recognition of sex-specific differences in carpal kinematics has various practical implications. With the knowledge that ‘column’-type wrists are associated with the type II lunate morphology^3^ and wrists with type II lunates are prone to degenerative cartilage changes at the proximal hamate^7^, one might infer an etiological relationship between the kinematic patterns seen primarily in women and proximal hamate osteoarthritis. Reporting on associations between sex and certain baseline kinematic patterns might aid in the surgical planning process by shifting the focus toward the restoration of baseline kinematics for a given patient. In the context of surgical treatments for chronic scapholunate instability, our results might caution against using procedures that limit flexion-extension of the scaphoid such as scapho-trapezio-trapezoid fusion^8^ for ‘column’-type wrists (i.e. women) and instead suggest the use of a procedure that better maintains scaphoid flexion-extension with wrist deviation in the coronal plane.

Our study had several limitations. The two-dimensional nature of the measurements and calculated metrics is a limiting factor when studying carpal kinematics, given that the carpal bones exhibit movement in several planes throughout various hand maneuvers^9^. The analysis of multiple points within the range of motion could add further knowledge regarding wrist kinematics, however this would require more x-ray acquisitions. This may be impractical in standard clinical settings, as these points will need to be pre-determined and tested, and would lead to additional radiation exposure to the patient. Our study therefore focused on utilizing the endpoints of the range of motion. Three-dimensional imaging modalities such as Magnetic Resonance Imaging^10,11^ and Computed Tomography^12,13^ have been used to investigate wrist kinematics and might allow us to better understand the complete trajectory of the scaphoid from radial to ulnar wrist deviation. Nevertheless, radiographs are easier to obtain, practical in clinical settings, correspond to lower ionizing radiation exposure than CT, and are less costly. While studies comparing radiography against such three-dimensional modalities are needed, our findings regarding the importance of stratifying by sex when evaluating scaphoid kinematics can help guide studies investigating carpal bone movement via any imaging modality. The ‘row-column’ theory of carpal kinematics describes a spectrum of kinematic patterns that are likely influenced by wrist ligament characteristics and bone morphologies. Future studies need to elucidate these associations for the implications of kinematic patterns to be understood.

Observation of scaphoid kinematic patterns also does not provide a comprehensive overview of kinematic patterns within the entire carpus due to the vast number of articulating joints and varying degrees of movement at each joint. Moreover, we were limited by the use of a single observer and the retrospective nature of our study limited our cohort size and ability to adjust for factors such as body frame size.

In conclusion, our study demonstrated sex-specific differences in the relationship between the degree of radial and ulnar wrist positioning and scaphoid positioning. Given that only men experience a change in the inclination of the scaphoid relative to the radius with wrist deviation in the coronal plane despite negligible changes in observed scaphoid length, we inferred that men have greater scaphoid movement in the coronal plane with radial and ulnar wrist deviation than women. In contrast, we did not observe significant differences in any of the scaphoid kinematic metrics for men versus women. This was consistent with previous studies^3,14^ and might suggest that examining the relationship between wrist deviation in the coronal plane and scaphoid characteristics can reveal more about sex-specific differences in kinematic patterns than comparing scaphoid characteristics on static images directly between the sexes.

## ACKNOWLEDGEMENTS

This study was funded by the National Institutes of Health grants 2K12 HD051958 and R03 EB015099. The views expressed in this article are the authors’ own and do not necessarily represent the views of the National Institutes of Health or the United States Government. The authors thank John Brock for help with materials for the manuscript.

**AUTHOR CONTRIBUTIONS STATEMENT**: All listed authors have made substantial contributions to research design, data analysis, and interpretation of data. Further, all authors have reviewed, provided comments on, and approved of the paper prior to submission for publication.

